# Optimal BR signalling is required for adequate cell wall orientation in the Arabidopsis root meristem

**DOI:** 10.1101/2021.03.02.433562

**Authors:** Zhenni Li, Ayala Sela, Yulia Fridman, Herman Höfte, Sigal Savaldi-Goldstein, Sebastian Wolf

## Abstract

The plant steroid hormones brassinosteroids (BRs) regulate growth in part through altering the properties of the cell wall, the extracellular matrix of plant cells. Conversely, cell wall signalling connects the state of cell wall homeostasis to the BR receptor complex and modulates BR activity. Here we report that both pectin-triggered cell wall signalling and impaired BR signalling result in altered cell wall orientation in the Arabidopsis root meristem. BR-induced defects in the orientation of newly placed walls are associated with aberrant localization of the cortical division zone but with normal specification of its positioning. Tissue-specific perturbations of BR signalling revealed that the cellular malfunction is unrelated to previously described whole organ growth defects. Thus, tissue type separates the pleiotropic effects of cell wall/BR signals and highlights their importance during cell wall placement.

## Introduction

The cell wall, a carbohydrate-rich structure surrounding all plant cells fulfils numerous essential functions in growth and development; it provides mechanical support, controls cellular adhesion and forms the interface with the environment (Cosgrove, 2016; Lampugnani et al., 2018; Voxeur and Hofte, 2016). Notably, since the cell wall prevents cell migration, tight control over the plane of cell wall deposition during cytokinesis is often assumed to be important for plants’ patterning and morphogenesis. However, whether there is a direct relationship between cell shape, in part controlled by cell division orientations, and organ shape is not clear (Beemster et al., 2003; Kaplan and Hagemann, 1991; Torres-Ruiz and Jurgens, 1994; Traas et al., 1995). Symmetric divisions add cells within tissues and asymmetric or formative divisions forms new tissue layers. During cytokinesis, the microtubules (MT) of the phragmoplast guide secretory traffic towards the cell plate, a radially expanding disk of membrane-engulfed cell wall material that gains in diameter until it fuses with the parental walls to conclude cell division (Livanos and Muller, 2019). Cell division plane orientation is pre-determined by the position of cortical division zone (CDZ). The MT network forms a transient ring structure known as preprophase band (PPB) at the periphery of the cell, marking the future fusion site of the cell plate and the parental wall during cytokinesis, and thus coincides with the orientation of the new cell wall (Livanos and Muller, 2019; Rasmussen and Bellinger, 2018; Rasmussen et al., 2013; Smith, 2001). While PPBs are at least partially dispensable for oriented cell division (Schaefer et al., 2017; Zhang et al., 2016), mutants in CDZ components like TAN1 and POK1,2 show severely altered division angles, suggesting that the phragmoplast lacks guidance in the absence of these factors (Lipka et al., 2014; Stockle et al., 2016; Walker et al., 2007). In light of the central role of cell wall biosynthesis during cytokinesis (Gu et al., 2016; Miart et al., 2014; Zuo et al., 2000) and a growing list of cell wall-mediated feedback signalling modules affecting a wide range of biological processes (Wolf, 2017), it is conceivable that cell wall state during mitosis/cytokinesis has to be monitored by cell wall surveillance systems (Rui and Dinneny, 2020; Vaahtera et al., 2019; Voxeur and Hofte, 2016).

Growth itself poses a threat to cell wall integrity and composition and physical properties of the cell wall have to be tightly monitored to ensure cell wall homeostasis and coordinated growth. For example, a compensatory response to cell wall challenge is mediated by RECEPTOR-LIKE PROTEIN 44 (RLP44) which is able to interact with the brassinosteroid (BR) receptor BRI1 (Holzwart et al., 2018) and its co-receptor BRI1-ASSOCIATED KINASE1 (BAK1) (Wolf et al., 2014) and promote BR signalling. More specifically, the degree of methylesterification (DM) of homogalacturonan (HG), a pectin component of the cell wall, has a profound impact on cell wall structure and mechanical properties: once secreted into the wall network, HG can be de-methylesterified by pectin methylesterases (PMEs), ubiquitous plant enzymes which are themselves regulated by PME inhibitor proteins (PMEIs). Reduction of PME activity through overexpression of *PECTIN METHYLESTERASE INHIBITOR 5* (*PMEI5*) activates the BR hormone signalling via RLP44, which in turn leads to a complex directional growth phenotype including organ fusion and root waving (Wolf et al., 2012; Wolf et al., 2014).

BR signalling has a context- and cell type-specific effects on plant growth and development (Ackerman-Lavert and Savaldi-Goldstein, 2019; Nolan et al., 2019; Planas-Riverola et al., 2019; Singh and Savaldi-Goldstein, 2015). In addition to the main receptor BRI1, BRs are also perceived by BRI1’s two close paralogs, BRL1 and BRL3 (Cano-Delgado et al., 2004; Zhou et al., 2004). *brl1* and *brl3* single and double mutants do not show growth defects and *bri1 brl1 brl3* triple mutant (*bri1-triple* from hereon) resembles *bri1*. However, the absence of *brl1* and *brl3* enhances *bri1’s* select vascular defects (Cano-Delgado et al., 2004; Holzwart et al., 2018; Kang et al., 2017). BR mutants have short meristems and reduced number of cell divisions (i.e. slow cell-cycle duration) in the longitudinal axis of the meristem (Cole et al., 2014; Gonzalez-Garcia et al., 2011; Hacham et al., 2011; Vragovic et al., 2015) and increased number of divisions, referred to as formative divisions, apparent in supernumerary cell files in root tissues (Holzwart et al., 2018; Kang et al., 2017). Tissue-specific BRI1 expression in the background of *bri1* and *bri1-triple* revealed differential effects of the receptor on organ growth. For example, epidermal (non-hair cell) BRI1 largely rescues bri1’s root length (Hacham et al., 2011). These roots have longer meristem as compared to the wild type, the cell count of which is is enhanced in *bri1-triple* and restrained by BRI1 in the stele (Vragovic et al., 2015). However, formative divisions are promoted by BRI1 in the stele and restrained by epidermal BRI1 (Kang et al., 2017). In addition, BRI1 activity in the protophloem cells of the vasculature can rescue the *bri-triple* root phenotype (Graeff et al., 2020; Kang et al., 2017). Together, tissue-specific approaches revealed that growth control by BRI1 involves cell autonomous and non-cell autonomous effects.

Here, we dissected the phenotype induced by PMEI5-mediated reduction in PME activity on the root apical meristem and revealed that pectin-triggered cell wall signalling leads to orientation defects of newly placed walls, which are dependent on BR signalling activation. These defects are independent of organ-level growth phenotypes and coincide with aberrant localization of the CDZ component POK1. Conversely, reduced BR signalling in receptors and biosynthetic mutants leads to cell shape defects similar to those observed in PMEIox, but unrelated to the enhanced formative division phenotypes. These cell shape defects are also separated from whole root growth defects. Thus, we reveal a role for cell wall and BR signalling in cell wall orientation.

## Results

### Pectin-triggered cell wall signalling leads to cell division defects that are RLP44 and BRI1 dependent

We have previously demonstrated that cell wall signalling connects the state of the cell wall to the BR pathway (Wolf et al., 2012). When de-methylesterified pectin becomes limiting, such as in PMEIox plants overexpressing *PMEI5*, elevated BR signalling counteracts loss of cell wall integrity and leads to BRI1-dependent macroscopic growth defects such as reduced root length and root waving ((Wolf et al., 2012); Figure 1A, Supplemental Figure 1A). To shed light on the cellular phenotype of PMEIox roots and to assess whether root waving is associated with meristematic defects, we analysed PMEIox root tips 5 days after germination (DAG) and found that the reduced root length of PMEIox was accompanied by an even stronger reduction in size and cell number of the RAM (Supplemental Figure 1A-C). Interestingly, we also revealed that in contrast to the stereotypical pattern of cellular morphology and tissue organization in wild type root tips ((Dolan et al., 1993), Figure 1B), PMEIox roots displayed a substantial amount of obliquely orientated transverse cell walls apparent in epidermis, cortex and endodermis (Figure 1C). This caused an irregular tissue organization and, in some severe cases, almost indiscernible tissue boundaries. We next asked whether these cell shape defects in PMEIox are also the result of elevated BR signalling mediated by RLP44 (Wolf et al., 2014). Consistent with this hypothesis, PMEIox suppressor mutants with lesions in *BRI1* (*cnu1*, (Wolf et al., 2012)) and *RLP44* (*cnu2*, (Wolf et al., 2014)), respectively, showed to a large extent normal cross wall orientations (Figure 1D, E) in line with the suppression of the macroscopic PMEIox defects, such a root waving and organ fusion (Figure 1A, (Wolf et al., 2012; Wolf et al., 2014). Moreover, as previously observed with the macroscopic phenotypes of PMEIox, expression of a GFP-tagged version of RLP44 under control of the CaMV35S promoter (RLP44-GFP) in *cnu2* resulted in oblique cell walls, indicating complementation (Supplemental Figure 2). Together, these results suggest that cell wall signalling–triggered elevation of BR signalling is causative for the oblique cell wall phenotype in PMEIox.

**Figure 1.**
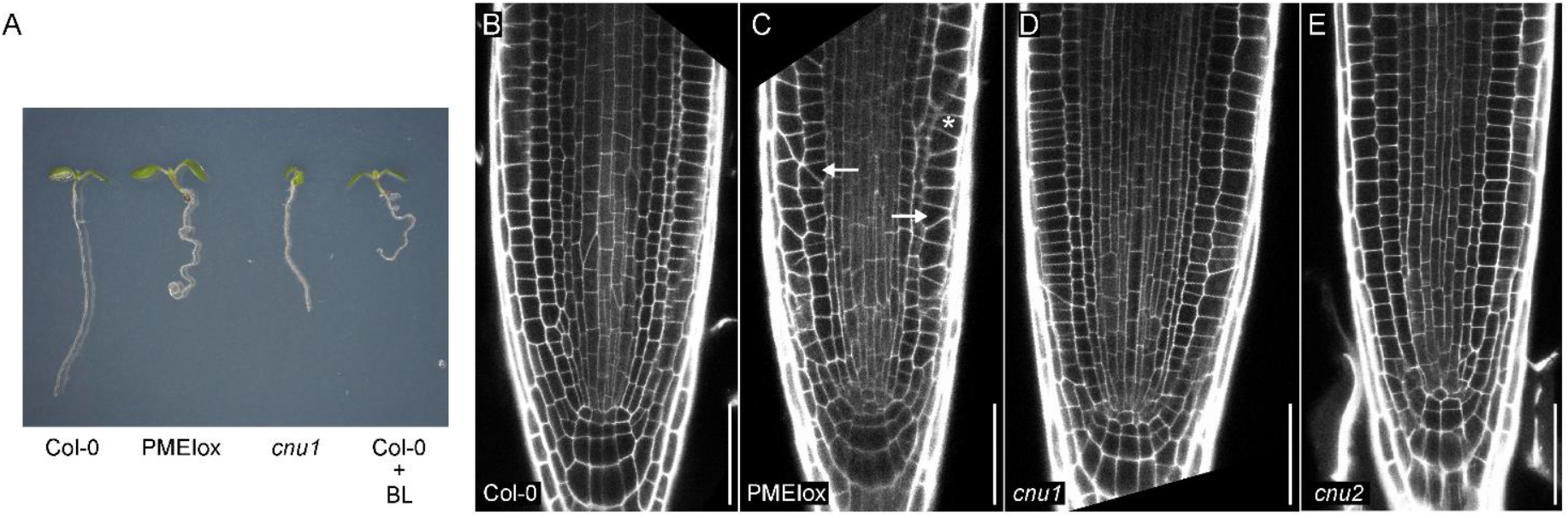
Cell wall signalling triggered by PMEIox alters root growth and cell wall orientation (A) PMEIox seedlings have a root waving phenotype caused by enhanced BR signalling. 5- day-old seedlings of Col-0, PMEIox, the PMEIox *bri1* suppressor mutant *cnu1*, and Col-0 seedlings grown on plates containing 5 nM of brassinolide (BL) are shown. (B-E). PMEIox plants show oblique cell walls in the root apical meristem dependent on BRI1 and RLP44. Cell division defects in PMEIox (C, arrows) are dependent on BRI1, mutated in the PMEIox suppressor mutant *cnu1* (D) and RLP44, mutated in the PMEIox suppressor mutant *cnu2* (E). Cells walls The root apical meristems are visualized using mPS-PI staining (Truernit et al., 2008). Bars = 50 µm.

### Cell wall perturbation in diverse cell types separates aberrant cell division, root waviness and root growth

Thus far, our results indicate that pectin-triggered cell wall signalling leads to root waving, root growth inhibition, and aberrant wall orientation that are BRI1-dependent. These pleiotropic phenotypes could be all linked or could result from unrelated processes. Since BR signalling is context-dependent, we employed a cell type-specific expression system to alter cell wall properties locally and followed the phenotypic consequences on the organ, tissue, and cellular level. We selected a number of tissue-specific promoters to drive expression of the chimeric transcription factor GR-LhG4 (Craft et al., 2005; Moore et al., 1998; Schurholz et al., 2018) in both cell types of the epidermis and in each of its cell types alone (i.e. hair cells and non-hair cells), cortex, endodermis, and xylem pole pericycle cells, complemented by ubiquitous expression (Supplemental Figure 3). GR-LhG4, in turn, triggers transcription of *PMEI5* under control of the synthetic pOp promoter in the presence of dexamethasone (DEX). Expression of *PMEI5* in the epidermis, driven by the *ML1* promoter (pML1>GR>PMEI5), or in hair cells (pCOBL9>GR>PMEI5) was sufficient to trigger root waving, similar to ubiquitous expression (pUBQ10>GR>PMEI5) (Figure 2A-C). In contrast, expression of *PMEI5* in non-hair cells, cortex, or differentiating endodermis cells did not affect directional root growth (Figure 2D-F). Interestingly, expression in the xylem pole pericycle cells of the stele (pXPP>GR>PMEI5) also led to root waving (Figure 2G). This suggests that cell wall-induced BR signalling in these cells causes organ-level responses, with the caveat that PMEI5 might be mobile in the cell wall. We then assessed the occurrence of oblique cell walls in the trans-activation lines. Notably, tissue-specific expression lines that displayed root waving, namely pML1>GR>PMEI5, pCOBL9>GR>PMEI5, and pXPP>GR>PMEI5, had normal cell wall angles in the RAM (Figure 3B-D), in contrast to ubiquitous *PMEI5* expression (Figure 3A), suggesting that root waving is independent from meristematic cell shape defects. Interestingly, trans-activation of *PMEI5* in cortex cells (pCO2>GR>PMEI5), which did not lead to root waving (Figure 2E), triggered PMEIox-like oblique cell walls cell autonomously in cortex cells and non-cell autonomously in epidermal cells (Figure 4). In addition, PMEI5 expression in cortex cells leads to a disrupted organization of the stem cell niche (Figure 4B). Moreover, and in contrast to PMEIox (Supplemental Figure 1), the profound effect on cell shape in pCO2>GR>PMEI5 plants was accompanied by only mild effects on root length, RAM size, and RAM cell number (Supplemental Figure 4). Taken together, these results reveal that triggering cell wall modification in either the epidermis or in XPP cells of the stele can lead to similar organ level responses and confirm that cell wall orientation alterations are independent of directional growth processes. In addition, meristematic cell shape defects did not affect organ level growth.

**Figure 2.**
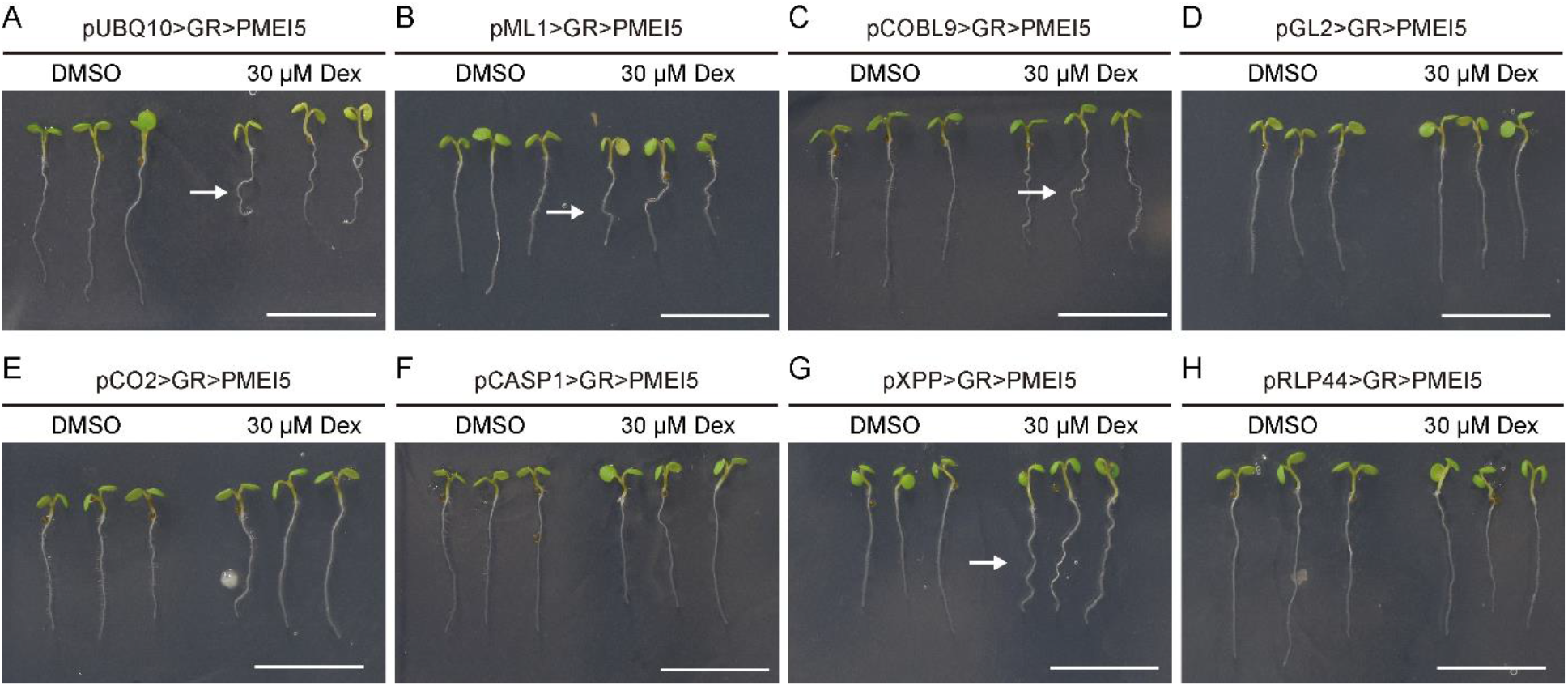
Cell wall perturbation in diverse cell types can lead to similar organ-level responses. Induction of PMEIox trans-activation ubiquitously (A), in the epidermis (B), hair cells (C), non- hair cells (D), meristematic cortex cells (E), differentiating endodermis (F), xylem pole pericycle cells (G), and in the RLP44 expression domain (H). Plants were germinated and grown on plates containing 30 µM Dexamethasone (Dex) or an equal volume of DMSO for five days. Bars = 1 cm.

**Figure 3.**
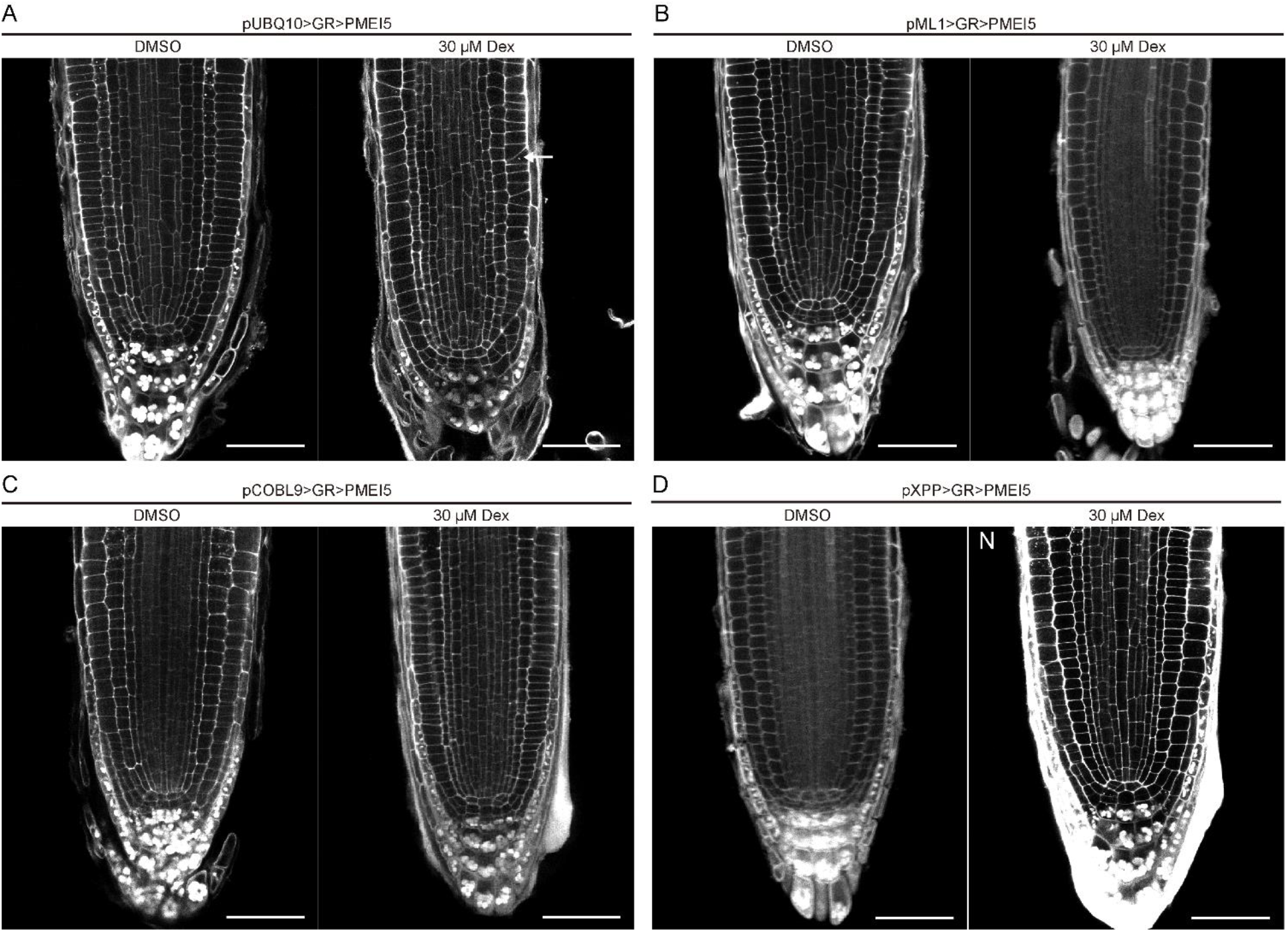
Tissue-specific expression of PMEIox reveals that root waving is independent from cell wall orientation defects. (A) Ubiquitous trans-activation of *PMEI5* recapitulates the PMEIox cell wall phenotype (arrow). (B-D) In lines with tissue-specific PMEI5 expression in epidermis, hair cells, or xylem pole pericycle cells, cell wall orientation in the RAM is normal. Bars = 50 µM. Cell walls are counterstained with mPS-PI.

**Figure 4.**
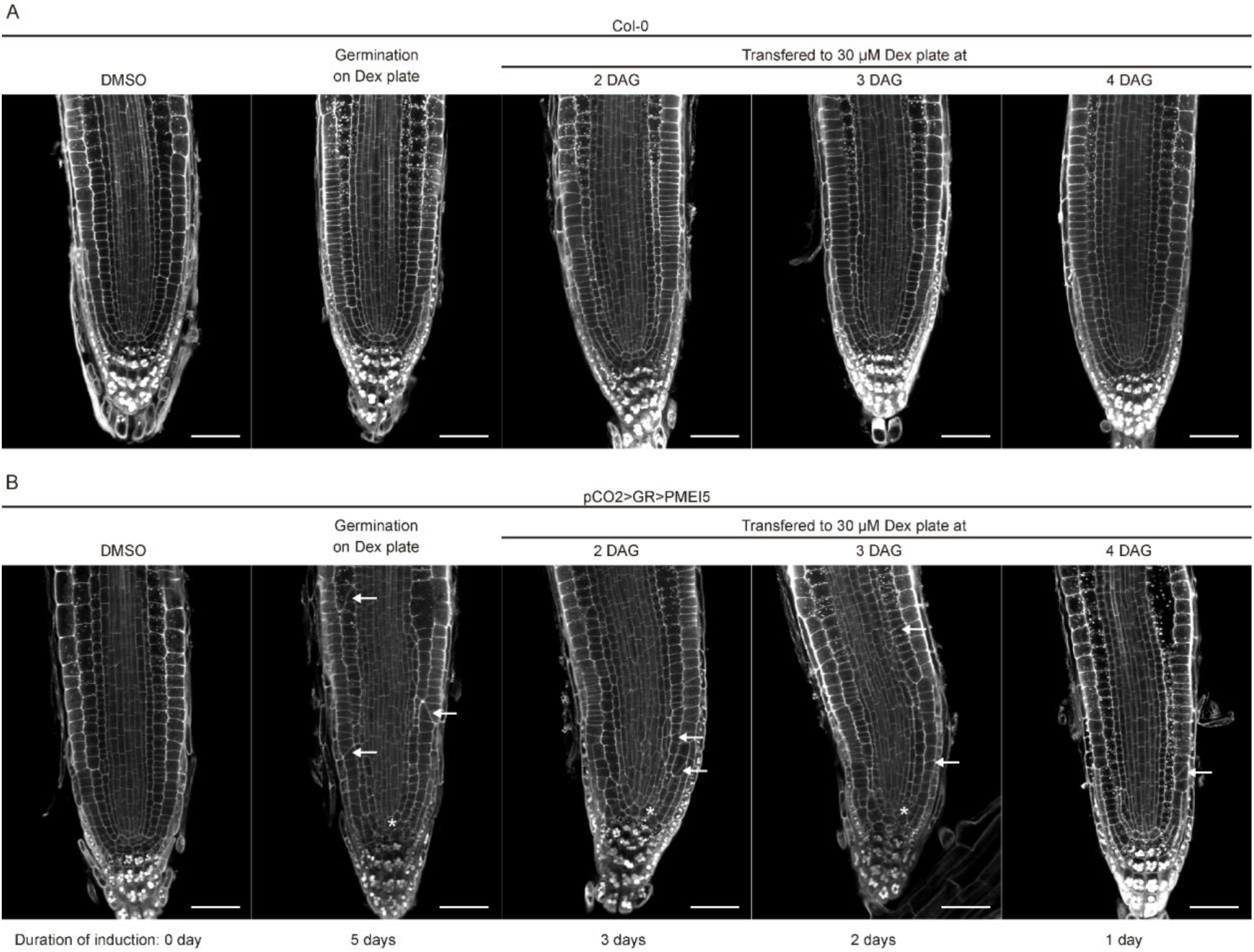
Induction of *PMEI5* expression in meristematic cortex cells leads to oblique cell walls in cortex and epidermis (arrows) and a disrupted stem cell region (asterisk). Plant were imaged at five days after germination on Dex-containing medium or after germination on DMSO- containing plates and transfer to induction medium at the indicated time points.

### Reduced BR signalling leads to an oblique newly wall orientation defects that can be separated from root growth

We next characterized the cross wall orientation in lines with reduced BR signalling and observed aberrant wall angles in *bri1* that were enhanced in *bri1-triple* mutants. The occurrence of oblique transversal walls within the cell files of the root meristem resulted in random perturbation of the cell files (Figure 5A), and thus unrelated to formative divisions (Holzwart et al., 2018; Kang et al., 2017). Together, optimal BR signalling strength is required for the maintenance of cell wall orientation (Figure 5). Quantification of aberrant cell walls in cortex cells revealed that the majority of *bri1-116* meristems had a least one oblique cortex cross wall per median confocal section, a phenotype which was enhanced in *bri-triple* mutants (Figure 5B). We recently reported that some functions of BRI1 are independent of classical BR signalling outputs (Holzwart et al., 2018; Holzwart et al., 2019). However, cell shape defects in *bri1* mutants seem to be caused by reduced BRI1/ligand dependent signalling, as the biosynthetic mutants *cpd* (Szekeres et al., 1996) and *det2* (Chory et al., 1991), as well as plants in which endogenous BRs were depleted by the application of brassinazole (BRZ) (Asami et al., 2000), displayed phenotypes comparable to *bri1* mutants (Figure 5B). To address the spatiotemporal impact of BR activity on the cell wall orientation, we used a previously reported collection of lines conferring tissue-specific BRI1 expression in *bri1* and *bri-triple* mutant backgrounds (Fridman et al., 2014; Hacham et al., 2011; Vragovic et al., 2015). Expression of BRI1 in non-hair cells (pGL2-BRI1) did not affect cortex cell wall orientation in the wild type, but slightly enhanced both the *bri1-116* and *bri-triple* phenotype (Figure 5C). In contrast, expression of BRI1 from the endodermis (pSCR-BRI1) or stele (pSHR-BRI1) largely rescued cell wall orientation in the cortex of *bri1-116* and *bri1-triple* (Figure 5C). Interestingly, the extent of BRI1’s effect on the aberrant wall angles did not correlate with its effect on meristem size and root length ((Hacham et al., 2011; Vragovic et al., 2015), Figure 5 D, E, Supplemental Figure 5). Specifically, GL2-BRI1 roots are longer than pSCR-BRI1 and pSHR-BRI1 and can have almost wild type length, in both *bri1* and *bri1-triple* backgrounds ((Hacham et al., 2011), Figure 5D, E). BRI1 activity in non-hair cells limits cell elongation in the elongation zone of the root, unless BRI1 in the neighbouring hair cells is also present (Fridman et al., 2014). High BRI1 in non-hair cells results in strong and mild waviness in a mutant and wild type background, respectively ((Fridman et al., 2014), Figure 5D). Thus, aberrant wall orientation is not correlated with root waviness. Together, as with PMEIox, the control of cell wall orientation is separated from other growth processes controlled by BRI1.

**Figure 5.**
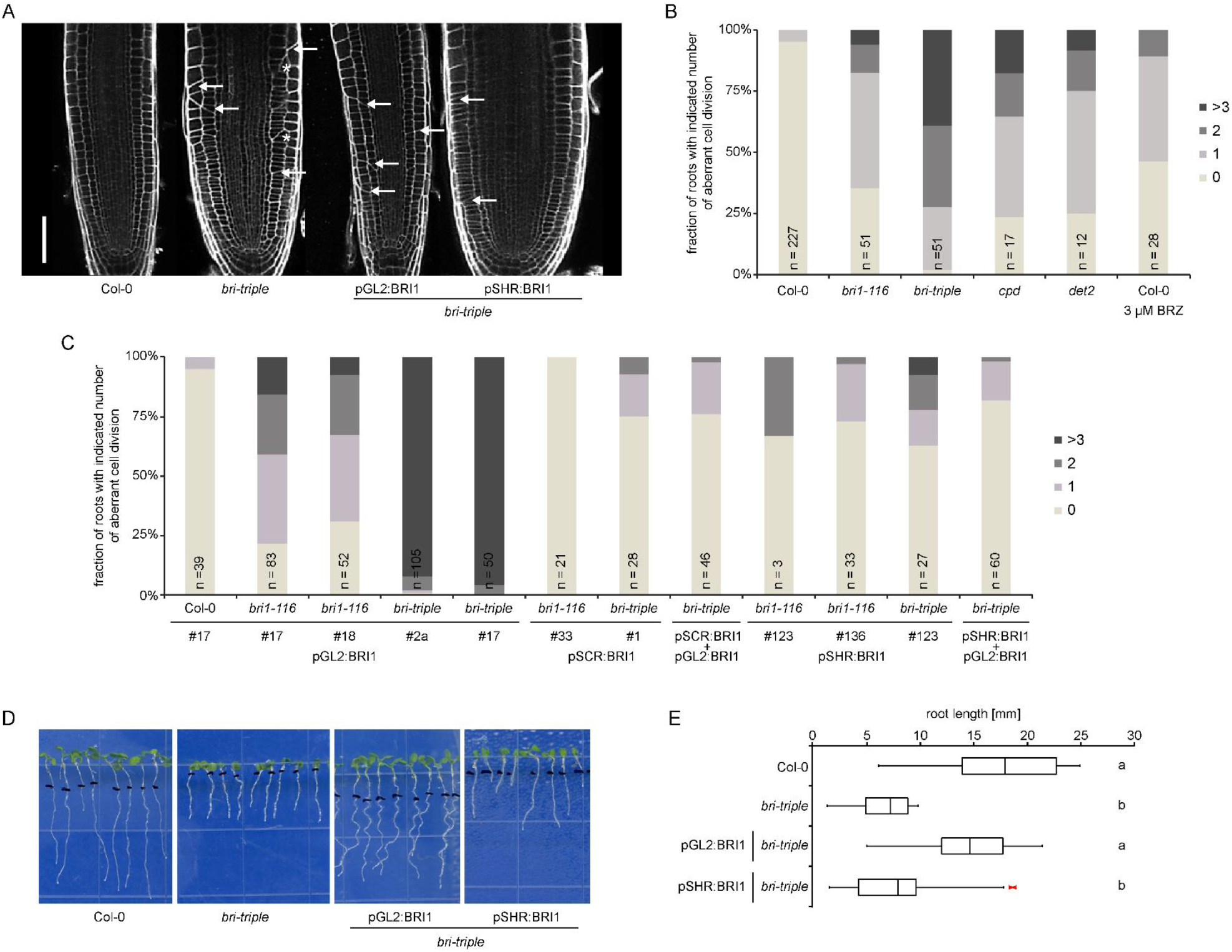
BR signalling is required for the maintenance of cell wall orientation. (A) Propidium Iodide-stained meristems of Col-0, *bri-triple* mutants and two *bri-triple* lines expressing BRI1 from cells of the epidermis (pGL2:BRI1) or stele (pSHR:BRI1). Note oblique transversal walls in epidermis, cortex, and endodermis (arrows), as well as random perturbation of cell files (asterisk). (B) Quantification of the fraction of cortex cells with oblique transversal walls in cortex cells from confocal section as in (A) for the indicated genotypes and wild type roots in which BRs were depleted by treatment with brassinazole. (C) Quantification of the fraction of cortex cells with altered cell shape of lines expressing BRI1 form the epidermis (pGL2-BRI1), endodermis (pSCR-BRI1), and stele (pSHR-BRI1) as well as their combinations in various backgrounds. (D) Growth phenotype of seven day old Col-0, *bri-triple*, and *bri-triple* plants expressing BRI1 from cells of the epidermis (pGL2:BRI1) or stele (pSHR:BRI1). (E) Quantification of root length as depicted in (D). n= 21-25. Letters indicate statistically significant difference after one factor ANOVA followed by Tukey’s post-hoc test.

### Cell wall perturbation by PMEIox leads to cytokinesis defects after specification of the CDZ

To test whether the aberrant cell wall placement in PMEIox was associated with defects in cell division plane orientation, we quantified the orientation of this plane at the different mitotic stages (Figure 6). The PPB transiently indicates positioning of the CDZ (Rasmussen et al., 2013). Hence, to determine whether cell wall perturbations affects this positioning, we used a fluorescently labelled tubulin (GFP-TUA6) that allows visualizing the PPB (Figure 6A and B). Quantitative analysis of PPB orientation in wild type cells revealed only minimal deviations from a position perpendicular to the cell long axis (<10°, Figure 6I). PMEIox displayed a very similar distribution, indicating that division plane orientation defects are independent of PPB positioning (Figure 6I). As the PPB disappears in pro-metaphase, we used a transgenic line expressing an RFP-fused Histone 2B to label chromatin and a GFP-fused plasma membrane protein Lti6b (Maizel et al., 2011) to visualize cell outlines and the forming cell plate. We quantified the orientation of the metaphase plate, the midline between sister chromatids, and the cell plate in metaphase, anaphase, and telophase, respectively. Interestingly, the orientation of both the Col-0 wild-type and PMEIox metaphase plates deviated considerably from a 90° angle relative to the cell long axis (Figure 6C, D, and J). A similar observation was made during anaphase (Figure 6E, F, and K). However, the forming cell plate during telophase was largely aligned with the expected division plane in Col-0, and deviation angles showed a very similar distribution to what was previously observed with the PPB (Figure 6G and L). In contrast, telophase cell plate orientation in PMEIox showed a distribution similar to what was observed in meta- and anaphase, suggesting defective re-alignment to the ∼90° angle observed in Col-0 (Figure 6H and L). In summary, the PPB orientation in wild type and PMEIox were indistinguishable from each other, suggesting that the oblique cell walls in PMEIox are not caused by MT misalignment. While both, wild type and PMEIox virtual division planes defined by the H2B marker, deviated markedly from the perpendicular PPB orientation during metaphase and anaphase, wild type cell plates during telophase aligned with the PPB position, whereas PMEIox cell plates did not.

**Figure 6.**
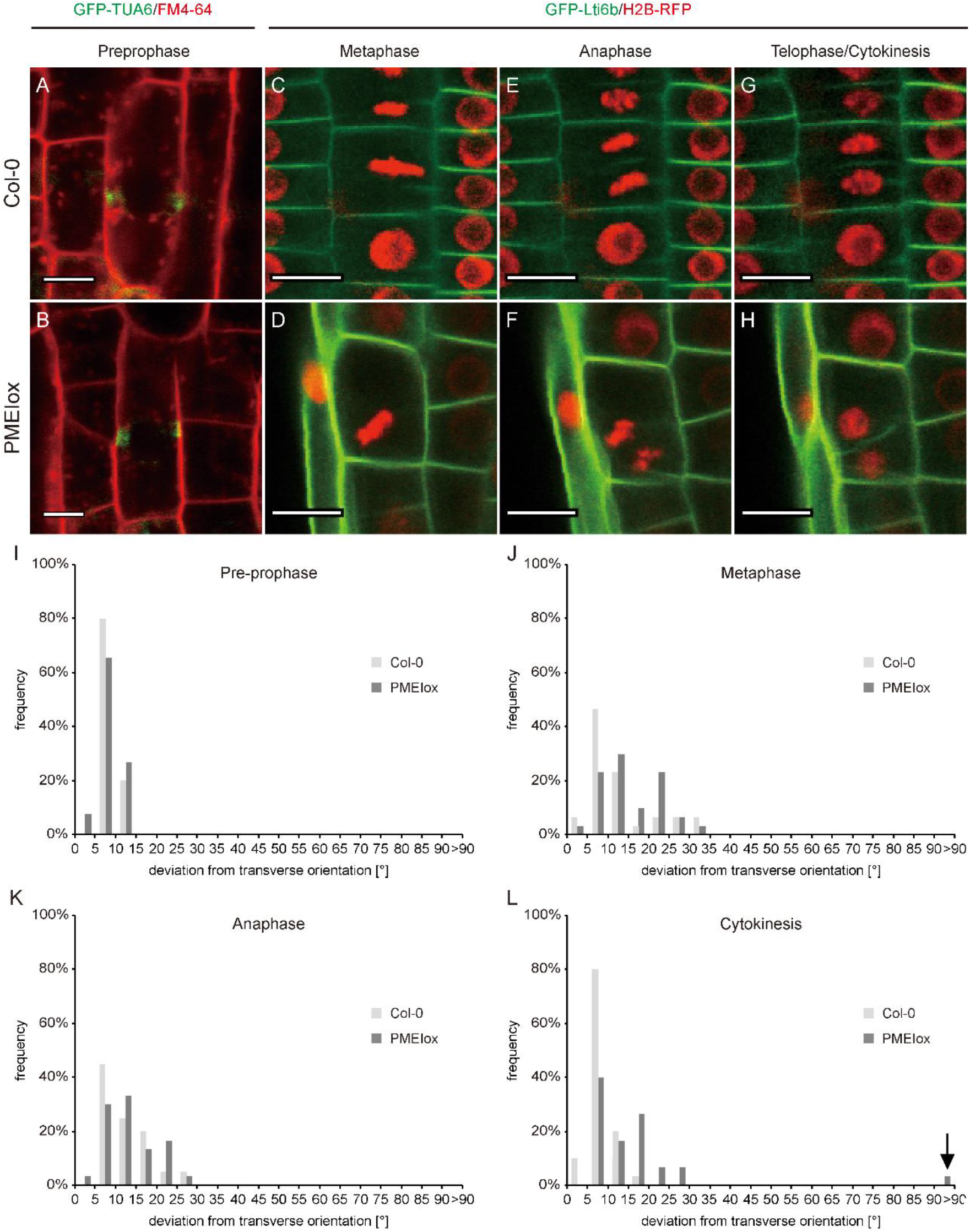
Cell wall perturbation by PMEIox leads to cell division defects after specification of the cortical division zone. (A, B) Orientation of pre-prophase bands labelled by GFP-TUA6 is transverse in both Col-0 (A) and PMEIox (B); see (I) for quantification. (C-F) Metaphase plate orientation (C, D; see J for quantification) and sister chromatid orientation during anaphase (E, F; see K for quantification) deviates from a 90°C angle in both the Col-0 wild type (C, E) and PMEIox (D, F). (G, H) Orientation of the cell plate in the Col-0 wild type stabilized at around 90 °C relative to the cell long axis (G), while PMEIox cell plates showed a wide range of orientations (H, see L for quantification). (I-L) Quantification of division plane orientation during mitosis and cytokinesis. Bars in histograms denote fraction of cells in bins bordered by angles indicated on the x-axis. n = 30 - 50. Bars = 20 µm in (A-H), = 10 µm in (I, J).

These observations are consistent with two hypotheses that could explain the aberrant cell wall angles in PMEIox. First, PMEIox cell plates might fail to find the CDZ initially marked by the PPB and occupied by POK1 and other components (Livanos and Muller, 2019). Second, the position of the cortical division zone might be mobile relative to the cell walls in PMEIox, while guidance is unaffected. To differentiate between the two possibilities, we introduced a fluorescently tagged version of POK1, YFP-POK1 (Lipka et al., 2014) into PMEIox and determined whether the cell plate fusion site coincided with the location of POK1 (Lipka et al., 2014; Muller et al., 2006). Cell plate fusion sites were marked by YFP-POK1, even when the cell plate fused at an angle deviating from 90° relative to the parental walls, suggesting that phragmoplast guidance towards CDZ components is unaffected in PMEIox (Figure 7). Instead, YFP-POK1, and thus presumably CDZ localization, seems to diverge from the position of the PPB. Together, cell plate fusion in PMEIox coincides with POK1 localization at aberrant positions, after specification of the CDZ. Our results show that optimal BR signalling strength is required to maintain the orientation of newly placed cell walls.

**Figure 7.**
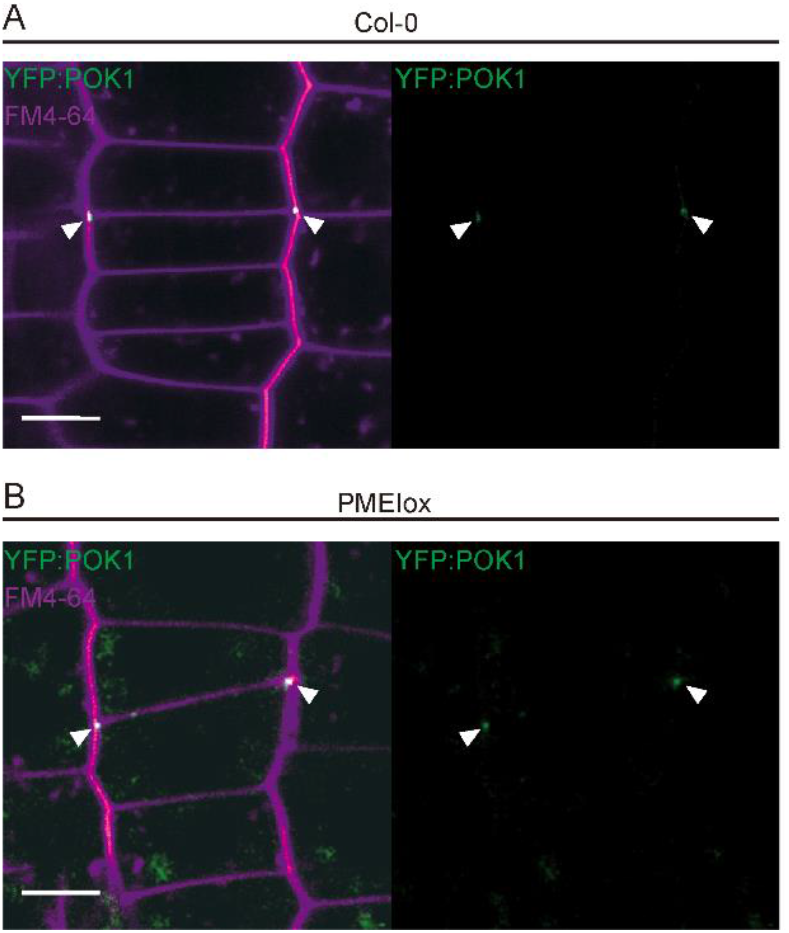
Cell plate fusion in PMEIox coincides with POK1 localization at aberrant positions. (A, B) Cell plate fusion site position and YFP-POK1 localization at the end of cytokinesis in Col-0 (A) and PMEIox (B). Note that the cell plate fusion at POK1-positive sites in the parental walls deviates from the normal position in PMEIox. White arrows heads indicated YPF-POK1- marked cell plate fusion sites. Membranes are labelled with FM4-64. Bars = 20 μm.

## Discussion

Here, we report that cell wall integrity and BR signalling are involved in the control of cell wall orientation in the Arabidopsis root. Both pectin-triggered cell wall signalling, as in PMEIox plants, and impaired BR signalling resulted in cross walls in the Arabidopsis root meristem. In PMEIox plants, pectin modification triggers an RLP44-mediated activation of BR signalling which, in turn, prevents loss of cell wall integrity, but results in a wide variety of growth related phenotypes ((Holzwart et al., 2018; Wolf et al., 2012; Wolf et al., 2014), this study). Aberrant cell wall orientation due to an altered cell wall in PMEIox seems to occur downstream of BR signalling, as mutation of *BRI1* or *RLP44* in the *cnu1* and *cnu2* mutants respectively, largely suppressed the oblique cell wall phenotype of PMEIox. Analysis of PMEIox cell divisions at the subcellular level revealed that the virtual cell division plane between meta- and telophase, marked by the midline between chromatin structures (metaphase plate and segregating sister chromatids), showed deviations from the 90° angle in both wild type and PMEIox cells. Wild type cell plates later returned to the previous position of the PPB, whereas PMEIox cell plates frequently did not. Our use of metaphase plate and sister chromatids as a read-out for the “virtual” cell division plane can be questioned as, for example, mitotic features like spindle orientation do not always correlate with division plane orientation (Cleary and Smith, 1998; Galatis et al., 1984; Marcus et al., 2003; Oud and Nanninga, 1992; Rasmussen et al., 2011; Rasmussen et al., 2013). However, that wild type cell division plane orientations show considerable variation and noise, but are later harmonized by interaction with CDZ components is in agreement with the phenotype of CDZ mutants such as *tan1, pok1pok2*, and *phgap1 phgap2* (Lipka et al., 2014; Muller et al., 2006; Smith et al., 1996; Stockle et al., 2016; Walker et al., 2007), in which oblique cell walls are presumably the result of a lack of phragmoplast guidance. Our analysis shows that PMEIox and WT behave similarly with respect to all aspects analysed up until the very last steps of cell division, during which PMEIox cell plates fail to re-align with the former orientation of the PPB. Notably, oblique cell walls in PMEIox seem to occur through a mechanism different from that in the aforementioned CDZ mutants, as the CDZ itself appears to have shifted from the position of the PPB, based on YFP-POK1 localization. This raises the question how the relative position of the CDZ is maintained in wild type cells and whether this could involve interactions with the cell wall as previously suggested (Smertenko et al., 2017). Supporting an involvement of the cell wall in CDZ maintenance is the observation that four way-junctions comprised by cross walls of adjacent cells fusing at a similar position of their shared longitudinal wall are actively avoided (Flanders et al., 1990; Martinez et al., 2018). PPBs are placed with an offset from the predicted division plane if a cross wall or PPB from an adjacent cell is found at this position and it is conceivable that the necessary signal to trigger this response involves the cell wall. In addition, cell wall attachment of the CDZ is a plausible way to maintain its relative position in the plasma membrane, in line with the observation that the cell wall is required for the maintenance of PIN polarity (Feraru et al., 2011) and for restricting the mobility of plasma membrane proteins in general (Martiniere et al., 2012; McKenna et al., 2019).

Whether there is a direct relationship between cell shape, in part controlled by cell division orientations and organ shape is a long standing question (Beemster et al., 2003; Hong et al., 2018; Kaplan and Hagemann, 1991; Martinez et al., 2017). Both, continuous pectin-triggered cell wall signaling activation and exogenous application of BRs results root growth alterations such as waviness and a reduction in root length ((Lanza et al., 2012; Wolf et al., 2012), this study). Here, we observed that cell type-specific induction of cell wall signaling separated root growth, its directionality, and cell wall orientation defects based on which cell type expressed PMEI5. Furthermore, pectin-triggered cell wall signaling and reduced BR signaling in *bri1* mutants led to similar cell wall orientation defects in the meristem, but only PMEIox roots showed directional growth phenotype at the organ level. In addition, plants expressing *PMEI5* in cortex cells showed pronounced cell division orientation defects without influencing organ morphology, growth, meristem size, or meristematic cell number, whereas epidermal expression of BRI1 in *bri1*-triple mutants rescued organ level growth but enhanced cell division orientation defects. These results demonstrate that aberrant cell shape are compatible with normal organ growth.

Reminiscent of previous findings demonstrating non-cell autonomous effects of BR signalling, cortical *PMEI5* expression led to oblique cell divisions in both epidermal and cortex cells. Unfortunately, potential mobility of the protein in the cell wall could not directly be addressed as all attempts to express a tagged version of PMEI5 failed, therefore we cannot determine whether the non-cell autonomous effect is directly linked to PMEI5 or to downstream signalling components. However, it is noteworthy that expression of *PMEI5* in two spatially separated cell types, hair cells in the epidermis and the XPP cells in the stele led to similar organ level responses (root waving). Future work needs to address how these organ level responses are connected to cellular effects of BR signalling and how the cell wall is connected to cell division orientation maintenance, taking into account potential mechanical feedbacks and the contribution of cell geometry and developmental signalling (Besson and Dumais, 2011; Besson and Dumais, 2014; Chakrabortty et al., 2018; Lintilhac and Vesecky, 1984; Louveaux et al., 2016; Martinez et al., 2018; Moukhtar et al., 2019; Yoshida et al., 2014).

The defects in cell wall orientation described here for BR receptor mutants appear random and it is unclear if these cell division defects are related to previously described disturbed cell files and altered tissue organization in rice *bri1* mutants (Nakamura et al., 2006). The use of a tissue-specific approach to perturb the BR signalling was an instrumental tool to disentangle the pleiotropic effects of the BR pathway. Hence, despite a ubiquitous expression of BRI1, it enabled to uncover tissue-specific effects on shoot growth, root meristem size, final cell size, root length and gene expression that were otherwise masked by alternative overexpression or loss-of-function studies (Singh and Savaldi-Goldstein, 2015). Here, we show that aberrant cell wall orientation can be largely rescued in both *bri1* and *bri1-triple* mutant backgrounds while other growth parameters are poorly rescued, and vice versa. For example, *pGL2-BRI1* in *bri1-triple* has long root meristems and almost wild type-like root length, but harbours severe wall orientation defects. However, *pSHR-BRI1* has a shorter root and wide meristem that largely rescued these defects. A tissue-specific approach also assisted in separating BRI1 control of phloem differentiation from that of growth (Graeff et al., 2020). Since BRI1 is not expressed in the phloem in the lines analysed here, restoration of BR signalling in diverse cell types is sufficient to control root length and orientation of transversal walls.

Taken together, our results demonstrate that cell wall integrity and optimal BR signaling levels are required for a correct cell wall placement. This control of cell wall orientation occurs both cell autonomously and non-cell autonomously and is uncoupled from organ level growth control.

## Materials and Methods

### Plant Material

All genotypes used in this study were in the Col-0 background and are listed in Table S1. For PMEox related experiments, seeds were sterilized with 1.3% (v/v) sodium hypochlorite (NaOCl) diluted in 70% ethanol for 3 minutes, then washed twice with 100% ethanol and dried in laminar flow hood. Seeds were sowed out on plate with growth medium containing half-strength (1⁄2) Murashige & Skroog (MS) medium (Duchefa), 1% D-sucrose (Car Roth) and 0.9% phytoagar (Duchefa) with pH adjusted to 5.8 with KOH. After 2 days stratification at 4°C in darkness, plates were placed vertically in long day conditions (16h light/ 8h dark cycles) with equal light conditions (approximately 100 μE m-2s-1) for 5 days. All analyses have been carried out on seedling of 5-dayold. For dexamethasone (Dex) induction on plate, desired amount of Dex (Signam-Aldrich #D4902) was added to the growth medium, equal volume of dimethyl sulfoxide (DMSO) has been added to the control plate. For Dex induction on soil, 30 μM Dex has been used to spray the aerial part of the plant and to water every other day starting from 3 days after transfer of seedling on soil. All plants are grown on soil under long day conditions (16h light/ 8h dark cycles) at 23°C with 65% humidity. For BRI1 related experiment (i.e. Figure 5), seeds were sterilized and grown as described in Fridman et al.

### Microscopy

Microscopic analyses have been carried out with Zeiss LSM 510 Meta, a Zeiss LSM710, and Leica TCS SP5 laser scanning confocal microscopes. For mTurquoise2, excitation wavelength of 458 nm was used and emission was collected between 460 and 520 nm. GFP was excited with a 488 nm laser line and fluorescence was collected between 490 and 530 nm, mVenus was excited with a 514 nm laser line and fluorescence was collected between 520 and 560 nm. For propidium iodide fluorescence, an excitation wavelength of 488 nm was used, whereas emission was collected between 600 and 670 nm. RFP and FM4-64 fluorescence was excited at 561 nm and emission was collected between 560 and 620 nm (RFP) or between 675 and 790 nm (FM4-64).

### Plasmid construction

All constructs were produced using GreenGate cloning (Lampropoulos et al., 2013) with modules and primers listed in Table S2. PCR products were generated using Q5® High-Fidelity DNA Polymerase (NEB) and column-purified by using GeneJET PCR Purification Kit (ThermoFisher), followed by restriction digest with *Eco*31I FD restriction enzyme (ThermoFisher) at 37°C for 15 minutes. Products were column-purified as described above. Empty entry vectors (pGGA000, pGGC000 (Lampropoulos et al., 2013) were digested and purified separately. Digested and purified insert and vector were ligated using Instant Sticky-end Ligase Master Mix (NEB) following instructions by the manufacturer. The ligation products were then used to transform chemically competent *E*.*coli* strain DH5*α* or XL1-blue and cultivated in LB medium supplied with Ampicillin. Plasmid sequences were verified by Sanger sequencing. Confirmed entry modules are ligated into intermediate vector by running GreenGate reaction (Lampropoulos et al., 2013). The generation of expression plasmids involved the creation of two intermediate constructs, one in pGGM000 carrying the GR-LhG4 expression cassette, and one in pGGN000, carrying the PMEI5 coding sequence under control of the pOp6 promoter. The assembly of two expression cassettes each carried by one intermediate vector was achieved by running the same GreenGate reaction and simply replacing the entry module and empty intermediate vector by intermediate module and empty pGGZ001 destination vector, respectively. Final plasmids were verified by colony PCR and restriction digest, and then used to transform *Agrobacterium tumefaciens* strain ASE (pSOUP+) carrying resistance to chloramphenicol, kanamycin and tetracycline. All constructs were transformed by the floral dip method as descried in (Clough and Bent, 1998; Zhang et al., 2006).

### mPS-PI staining

Staining of cell outline with propidium iodide (PI, Sigma-Aldrich #P4170) with modified pseudo-PI satining method has been performed as described in (Truernit et al., 2008) with modifications. Seedlings at 5 DAG were fixed in solution containing 50% methanol and 10% acetic acid for 3 days at 4 °C. Samples were then washed twice with H_2_O and incubated in 1% periodic acid (Sigma-Aldrich #P0430) at room temperature for 40 minutes. Samples were washed twice with H_2_O and then stained with 100 μg/mL PI freshly diluted in Schiff’s reagent (100 mM sodium metabisulphite, 75 mM HCl). Stained samples were transferred onto microscope slides covered by chlorohydrate solution (4 g chloral hydrate, 1 mL glycerol, 2 mL H_2_O) and incubated overnight at room temperature in a closed environment. Excess of chlorohydrate solution was removed and several drops of Hoyer’s solution (3 g gum arabic, 20 g chloral hydrate, 2 g glycerol, 5 mL H_2_O) was added to the samples, which were at the end covered gently by cover splits and stayed at room temperature for 3 days before imaging. Orientation of mitotic structures, RAM length, and meristematic cell number were measured using ImageJ. The meristem was defined as the region between the quiescent centre and the first cortex cells with twice the length of the previous cell.

## Acknowledgements

The authors thank Sabine Müller for the YFP-POK1 line.

## Summary Statement

Both increased and reduced BR signalling strength results in altered cell wall orientation in the Arabidopsis root, uncoupled from whole-root growth control.

## References

Ackerman-Lavert, M. and Savaldi-Goldstein, S. (2019). Growth models from a brassinosteroid perspective. Curr Opin Plant Biol 53, 90–97.

Asami, T., Min, Y. K., Nagata, N., Yamagishi, K., Takatsuto, S., Fujioka, S., Murofushi, N., Yamaguchi, I. and Yoshida, S. (2000). Characterization of brassinazole, a triazole-type brassinosteroid biosynthesis inhibitor. Plant Physiol 123, 93–100.

Beemster, G. T., Fiorani, F. and Inze, D. (2003). Cell cycle: the key to plant growth control? Trends Plant Sci 8, 154–158.

Besson, S. and Dumais, J. (2011). Universal rule for the symmetric division of plant cells. Proc Natl Acad Sci U S A 108, 6294–6299.

————(2014). Stochasticity in the symmetric division of plant cells: when the exceptions are the rule. Frontiers in plant science 5, 538.

Cano-Delgado, A., Yin, Y., Yu, C., Vafeados, D., Mora-Garcia, S., Cheng, J. C., Nam, K. H., Li, J. and Chory, J. (2004). BRL1 and BRL3 are novel brassinosteroid receptors that function in vascular differentiation in Arabidopsis. Development 131, 5341–5351.

Chakrabortty, B., Willemsen, V., de Zeeuw, T., Liao, C. Y., Weijers, D., Mulder, B. and Scheres, B. (2018). A Plausible Microtubule-Based Mechanism for Cell Division Orientation in Plant Embryogenesis. Curr Biol 28, 3031–3043 e3032.

Chory, J., Nagpal, P. and Peto, C. A. (1991). Phenotypic and Genetic Analysis of det2, a New Mutant That Affects Light-Regulated Seedling Development in Arabidopsis. Plant Cell 3, 445–459.

Cleary, A. L. and Smith, L. G. (1998). The Tangled1 gene is required for spatial control of cytoskeletal arrays associated with cell division during maize leaf development. Plant Cell 10, 1875–1888.

Clough, S. J. and Bent, A. F. (1998). Floral dip: a simplified method for Agrobacterium-mediated transformation of Arabidopsis thaliana. Plant J 16, 735–743.

Cole, R. A., McInally, S. A. and Fowler, J. E. (2014). Developmentally distinct activities of the exocyst enable rapid cell elongation and determine meristem size during primary root growth in Arabidopsis. BMC Plant Biol 14, 386.

Cosgrove, D. J. (2016). Plant cell wall extensibility: connecting plant cell growth with cell wall structure, mechanics, and the action of wall-modifying enzymes. J Exp Bot 67, 463–476.

Craft, J., Samalova, M., Baroux, C., Townley, H., Martinez, A., Jepson, I., Tsiantis, M. and Moore, I. (2005). New pOp/LhG4 vectors for stringent glucocorticoid-dependent transgene expression in Arabidopsis. Plant J 41, 899–918.

Dolan, L., Janmaat, K., Willemsen, V., Linstead, P., Poethig, S., Roberts, K. and Scheres, B. (1993). Cellular organisation of the Arabidopsis thaliana root. Development 119, 71–84.

Feraru, E., Feraru, M. I., Kleine-Vehn, J., Martiniere, A., Mouille, G., Vanneste, S., Vernhettes, S., Runions, J. and Friml, J. (2011). PIN polarity maintenance by the cell wall in Arabidopsis. Curr Biol 21, 338–343.

Flanders, D. J., Rawlins, D. J., Shaw, P. J. and Lloyd, C. W. (1990). Nucleus-associated microtubules help determine the division plane of plant epidermal cells: avoidance of four-way junctions and the role of cell geometry. J Cell Biol 110, 1111–1122.

Fridman, Y., Elkouby, L., Holland, N., Vragovic, K., Elbaum, R. and Savaldi-Goldstein, S. (2014). Root growth is modulated by differential hormonal sensitivity in neighboring cells. Genes Dev 28, 912–920.

Galatis, B., Apostolakos, P. and Katsaros, C. (1984). Positional inconsistency between preprophase microtubule band and final cell plate arrangement during triangular subsidiary cell and atypical hair cell formation in two Triticum species. Canadian Journal of Botany 62, 343–359.

Gonzalez-Garcia, M. P., Vilarrasa-Blasi, J., Zhiponova, M., Divol, F., Mora-Garcia, S., Russinova, E. and Cano-Delgado, A. I. (2011). Brassinosteroids control meristem size by promoting cell cycle progression in Arabidopsis roots. Development 138, 849–859.

Graeff, M., Rana, S., Marhava, P., Moret, B. and Hardtke, C. S. (2020). Local and Systemic Effects of Brassinosteroid Perception in Developing Phloem. Curr Biol 30, 1626–1638 e1623.

Gu, F., Bringmann, M., Combs, J. R., Yang, J., Bergmann, D. C. and Nielsen, E. (2016). Arabidopsis CSLD5 Functions in Cell Plate Formation in a Cell Cycle-Dependent Manner. Plant Cell 28, 1722–1737.

Hacham, Y., Holland, N., Butterfield, C., Ubeda-Tomas, S., Bennett, M. J., Chory, J. and Savaldi-Goldstein, S. (2011). Brassinosteroid perception in the epidermis controls root meristem size. Development 138, 839–848.

Holzwart, E., Huerta, A. I., Glockner, N., Garnelo Gomez, B., Wanke, F., Augustin, S., Askani, J. C., Schurholz, A. K., Harter, K. and Wolf, S. (2018). BRI1 controls vascular cell fate in the Arabidopsis root through RLP44 and phytosulfokine signaling. Proc Natl Acad Sci U S A.

Holzwart, E., Wanke, F., Glockner, N., Hofte, H., Harter, K. and Wolf, S. (2019). A mutant allele uncouples the brassinosteroid-dependent and independent functions of BRASSINOSTEROID INSENSITIVE 1. Plant Physiol.

Hong, L., Dumond, M., Zhu, M., Tsugawa, S., Li, C. B., Boudaoud, A., Hamant, O. and Roeder, A. H. K. (2018). Heterogeneity and Robustness in Plant Morphogenesis: From Cells to Organs. Annu Rev Plant Biol 69, 469–495.

Kang, Y. H., Breda, A. and Hardtke, C. S. (2017). Brassinosteroid signaling directs formative cell divisions and protophloem differentiation in Arabidopsis root meristems. Development 144, 272–280.

Kaplan, D. R. and Hagemann, W. (1991). The relationship of cell and organism in vascular plants. Bioscience 41, 693–703.

Lampropoulos, A., Sutikovic, Z., Wenzl, C., Maegele, I., Lohmann, J. U. and Forner, J. (2013). GreenGate---a novel, versatile, and efficient cloning system for plant transgenesis. PLoS One 8, e83043.

Lampugnani, E. R., Khan, G. A., Somssich, M. and Persson, S. (2018). Building a plant cell wall at a glance. J Cell Sci 131.

Lanza, M., Garcia-Ponce, B., Castrillo, G., Catarecha, P., Sauer, M., Rodriguez-Serrano, M., Paez-Garcia, A., Sanchez-Bermejo, E., T, C. M., Leo del Puerto, Y., et al. (2012). Role of actin cytoskeleton in brassinosteroid signaling and in its integration with the auxin response in plants. Dev Cell 22, 1275–1285.

Lintilhac, P. M. and Vesecky, T. B. (1984). Stress-induced alignment of division plane in plant tissues grown in vitro. Nature 307, 363–364.

Lipka, E., Gadeyne, A., Stockle, D., Zimmermann, S., De Jaeger, G., Ehrhardt, D. W., Kirik, V., Van Damme, D. and Muller, S. (2014). The Phragmoplast-Orienting Kinesin-12 Class Proteins Translate the Positional Information of the Preprophase Band to Establish the Cortical Division Zone in Arabidopsis thaliana. Plant Cell 26, 2617–2632.

Livanos, P. and Muller, S. (2019). Division Plane Establishment and Cytokinesis. Annu Rev Plant Biol 70, 239–267.

Louveaux, M., Julien, J. D., Mirabet, V., Boudaoud, A. and Hamant, O. (2016). Cell division plane orientation based on tensile stress in Arabidopsis thaliana. Proc Natl Acad Sci U S A 113, E4294–4303.

Maizel, A., von Wangenheim, D., Federici, F., Haseloff, J. and Stelzer, E. H. (2011). High-resolution live imaging of plant growth in near physiological bright conditions using light sheet fluorescence microscopy. Plant J 68, 377–385.

Marcus, A. I., Li, W., Ma, H. and Cyr, R. J. (2003). A kinesin mutant with an atypical bipolar spindle undergoes normal mitosis. Mol Biol Cell 14, 1717–1726.

Martinez, P., Allsman, L. A., Brakke, K. A., Hoyt, C., Hayes, J., Liang, H., Neher, W., Rui, Y., Roberts, A. M., Moradifam, A., et al. (2018). Predicting Division Planes of Three-Dimensional Cells by Soap-Film Minimization. Plant Cell 30, 2255–2266.

Martinez, P., Luo, A., Sylvester, A. and Rasmussen, C. G. (2017). Proper division plane orientation and mitotic progression together allow normal growth of maize. Proc Natl Acad Sci U S A 114, 2759–2764.

Martiniere, A., Lavagi, I., Nageswaran, G., Rolfe, D. J., Maneta-Peyret, L., Luu, D. T., Botchway, S. W., Webb, S. E., Mongrand, S., Maurel, C., et al. (2012). Cell wall constrains lateral diffusion of plant plasma-membrane proteins. Proc Natl Acad Sci U S A 109, 12805–12810.

McKenna, J. F., Rolfe, D. J., Webb, S. E. D., Tolmie, A. F., Botchway, S. W., Martin-Fernandez, M. L., Hawes, C. and Runions, J. (2019). The cell wall regulates dynamics and size of plasma-membrane nanodomains in Arabidopsis. Proc Natl Acad Sci U S A.

Miart, F., Desprez, T., Biot, E., Morin, H., Belcram, K., Hofte, H., Gonneau, M. and Vernhettes, S. (2014). Spatio-temporal analysis of cellulose synthesis during cell plate formation in Arabidopsis. Plant J 77, 71–84.

Moore, I., Galweiler, L., Grosskopf, D., Schell, J. and Palme, K. (1998). A transcription activation system for regulated gene expression in transgenic plants. Proc Natl Acad Sci U S A 95, 376–381.

Moukhtar, J., Trubuil, A., Belcram, K., Legland, D., Khadir, Z., Urbain, A., Palauqui, J. C. and Andrey, P. (2019). Cell geometry determines symmetric and asymmetric division plane selection in Arabidopsis early embryos. PLoS computational biology 15, e1006771.

Muller, S., Han, S. and Smith, L. G. (2006). Two kinesins are involved in the spatial control of cytokinesis in Arabidopsis thaliana. Curr Biol 16, 888–894.

Nakamura, A., Fujioka, S., Sunohara, H., Kamiya, N., Hong, Z., Inukai, Y., Miura, K., Takatsuto, S., Yoshida, S., Ueguchi-Tanaka, M., et al. (2006). The role of OsBRI1 and its homologous genes, OsBRL1 and OsBRL3, in rice. Plant Physiol 140, 580–590.

Nolan, T., Vukasinovic, N., Liu, D., Russinova, E. and Yin, Y. (2019). Brassinosteroids: Multi-Dimensional Regulators of Plant Growth, Development, and Stress Responses. Plant Cell.

Oud, J. L. and Nanninga, N. (1992). Cell shape, chromosome orientation and the position of the plane of division in Vicia faba root cortex cells. Journal of Cell Science 103, 847.

Planas-Riverola, A., Gupta, A., Betegon-Putze, I., Bosch, N., Ibanes, M. and Cano-Delgado, A. I. (2019). Brassinosteroid signaling in plant development and adaptation to stress. Development 146.

Rasmussen, C. G. and Bellinger, M. (2018). An overview of plant division-plane orientation. New Phytol 219, 505–512.

Rasmussen, C. G., Sun, B. and Smith, L. G. (2011). Tangled localization at the cortical division site of plant cells occurs by several mechanisms. J Cell Sci 124, 270–279.

Rasmussen, C. G., Wright, A. J. and Muller, S. (2013). The role of the cytoskeleton and associated proteins in determination of the plant cell division plane. Plant J 75, 258–269.

Rui, Y. and Dinneny, J. R. (2020). A wall with integrity: surveillance and maintenance of the plant cell wall under stress. New Phytol 225, 1428–1439.

Schaefer, E., Belcram, K., Uyttewaal, M., Duroc, Y., Goussot, M., Legland, D., Laruelle, E., de Tauzia-Moreau, M. L., Pastuglia, M. and Bouchez, D. (2017). The preprophase band of microtubules controls the robustness of division orientation in plants. Science 356, 186–189.

Schurholz, A. K., Lopez-Salmeron, V., Li, Z., Forner, J., Wenzl, C., Gaillochet, C., Augustin, S., Barro, A. V., Fuchs, M., Gebert, M., et al. (2018). A Comprehensive Toolkit for Inducible, Cell Type-Specific Gene Expression in Arabidopsis. Plant Physiol 178, 40–53.

Singh, A. P. and Savaldi-Goldstein, S. (2015). Growth control: brassinosteroid activity gets context. J Exp Bot 66, 1123–1132.

Smertenko, A., Assaad, F., Baluska, F., Bezanilla, M., Buschmann, H., Drakakaki, G., Hauser, M. T., Janson, M., Mineyuki, Y., Moore, I., et al. (2017). Plant Cytokinesis: Terminology for Structures and Processes. Trends Cell Biol 27, 885–894.

Smith, L. G. (2001). Plant cell division: building walls in the right places. Nat Rev Mol Cell Biol 2, 33–39.

Smith, L. G., Hake, S. and Sylvester, A. W. (1996). The tangled-1 mutation alters cell division orientations throughout maize leaf development without altering leaf shape. Development 122, 481–489.

Stockle, D., Herrmann, A., Lipka, E., Lauster, T., Gavidia, R., Zimmermann, S. and Muller, S. (2016). Putative RopGAPs impact division plane selection and interact with kinesin-12 POK1. Nat Plants 2, 16120.

Szekeres, M., Nemeth, K., Koncz-Kalman, Z., Mathur, J., Kauschmann, A., Altmann, T., Redei, G. P., Nagy, F., Schell, J. and Koncz, C. (1996). Brassinosteroids rescue the deficiency of CYP90, a cytochrome P450, controlling cell elongation and de-etiolation in Arabidopsis. Cell 85, 171–182.

Torres-Ruiz, R. A. and Jurgens, G. (1994). Mutations in the FASS gene uncouple pattern formation and morphogenesis in Arabidopsis development. Development 120, 2967–2978.

Traas, J., Bellini, C., Nacry, P., Kronenberger, J., Bouchez, D. and Caboche, M. (1995). Normal differentiation patterns in plants lacking microtubular preprophase bands. Nature 375, 676–677.

Truernit, E., Bauby, H., Dubreucq, B., Grandjean, O., Runions, J., Barthelemy, J. and Palauqui, J. C. (2008). High-resolution whole-mount imaging of three-dimensional tissue organization and gene expression enables the study of Phloem development and structure in Arabidopsis. Plant Cell 20, 1494–1503.

Vaahtera, L., Schulz, J. and Hamann, T. (2019). Cell wall integrity maintenance during plant development and interaction with the environment. Nat Plants 5, 924–932.

Voxeur, A. and Hofte, H. (2016). Cell wall integrity signaling in plants: “To grow or not to grow that’s the question”. Glycobiology.

Vragovic, K., Sela, A., Friedlander-Shani, L., Fridman, Y., Hacham, Y., Holland, N., Bartom, E., Mockler, T. C. and Savaldi-Goldstein, S. (2015). Translatome analyses capture of opposing tissue-specific brassinosteroid signals orchestrating root meristem differentiation. Proc Natl Acad Sci U S A 112, 923–928.

Walker, K. L., Muller, S., Moss, D., Ehrhardt, D. W. and Smith, L. G. (2007). Arabidopsis TANGLED identifies the division plane throughout mitosis and cytokinesis. Curr Biol 17, 1827–1836.

Wolf, S. (2017). Plant cell wall signalling and receptor-like kinases. The Biochemical journal 474, 471–492.

Wolf, S., Mravec, J., Greiner, S., Mouille, G. and Hofte, H. (2012). Plant cell wall homeostasis is mediated by brassinosteroid feedback signaling. Curr Biol 22, 1732–1737.

Wolf, S., van der Does, D., Ladwig, F., Sticht, C., Kolbeck, A., Schurholz, A. K., Augustin, S., Keinath, N., Rausch, T., Greiner, S., et al. (2014). A receptor-like protein mediates the response to pectin modification by activating brassinosteroid signaling. Proc Natl Acad Sci U S A.

Yoshida, S., Barbier de Reuille, P., Lane, B., Bassel, G. W., Prusinkiewicz, P., Smith, R. S. and Weijers, D. (2014). Genetic control of plant development by overriding a geometric division rule. Dev Cell 29, 75–87.

Zhang, X., Henriques, R., Lin, S. S., Niu, Q. W. and Chua, N. H. (2006). Agrobacterium-mediated transformation of Arabidopsis thaliana using the floral dip method. Nature protocols 1, 641–646.

Zhang, Y., Iakovidis, M. and Costa, S. (2016). Control of patterns of symmetric cell division in the epidermal and cortical tissues of the Arabidopsis root. Development 143, 978–982.

Zhou, A., Wang, H., Walker, J. C. and Li, J. (2004). BRL1, a leucine-rich repeat receptor-like protein kinase, is functionally redundant with BRI1 in regulating Arabidopsis brassinosteroid signaling. Plant J 40, 399–409.

Zuo, J., Niu, Q. W., Nishizawa, N., Wu, Y., Kost, B. and Chua, N. H. (2000). KORRIGAN, an Arabidopsis endo-1,4-beta-glucanase, localizes to the cell plate by polarized targeting and is essential for cytokinesis. Plant Cell 12, 1137–1152.

