## Supplemental Figures for "Optimal BR signalling is required for adequate cell wall orientation in the Arabidopsis root meristem"

Suppplementary Figures

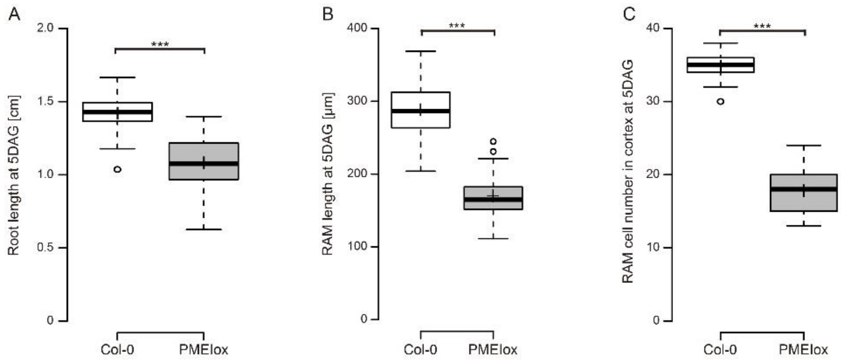

Supplemental Figure 1. PMEIox plants show reduced root length (A), RAM length (B) and RAM cell number five days after germination (5DAG). Asterisk indicates a statistically significant difference as determined by a two-tail t-test with $p < 0.001$. n = 34-46. Boxes indicate median, upper and lower quartile, whiskers indicate minimum and maximum except outliers beyond 1.5x interquartile range, which are indicated as dots.

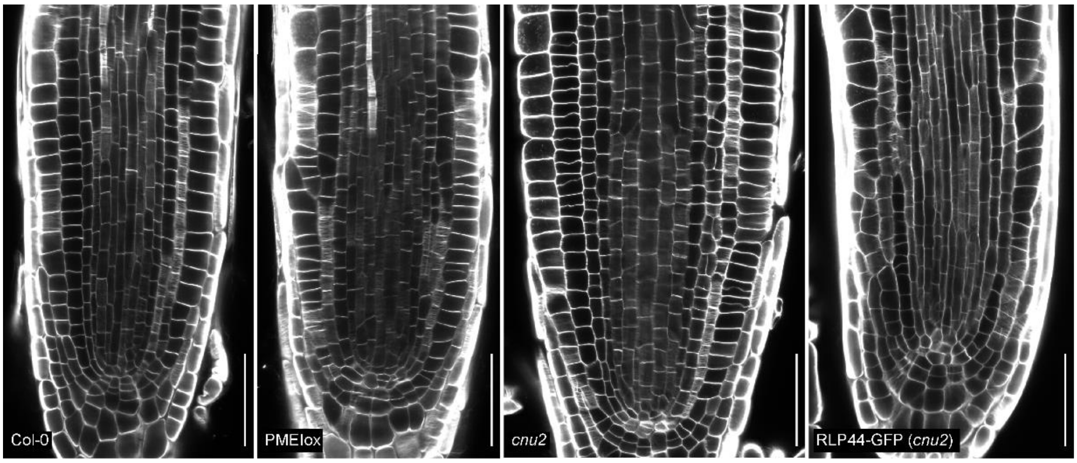

Supplemental Figure 2. Expression of RLP44-GFP in *cnu2* mutants restores aberrant cell division plane orientation. Median sections through root tips of Col-0, PMEIox, the PMEIox suppressor mutant *cnu2* and a complemented line expressing RLP44-GFP in the *cnu2* background. Scale bars = 50 µm.

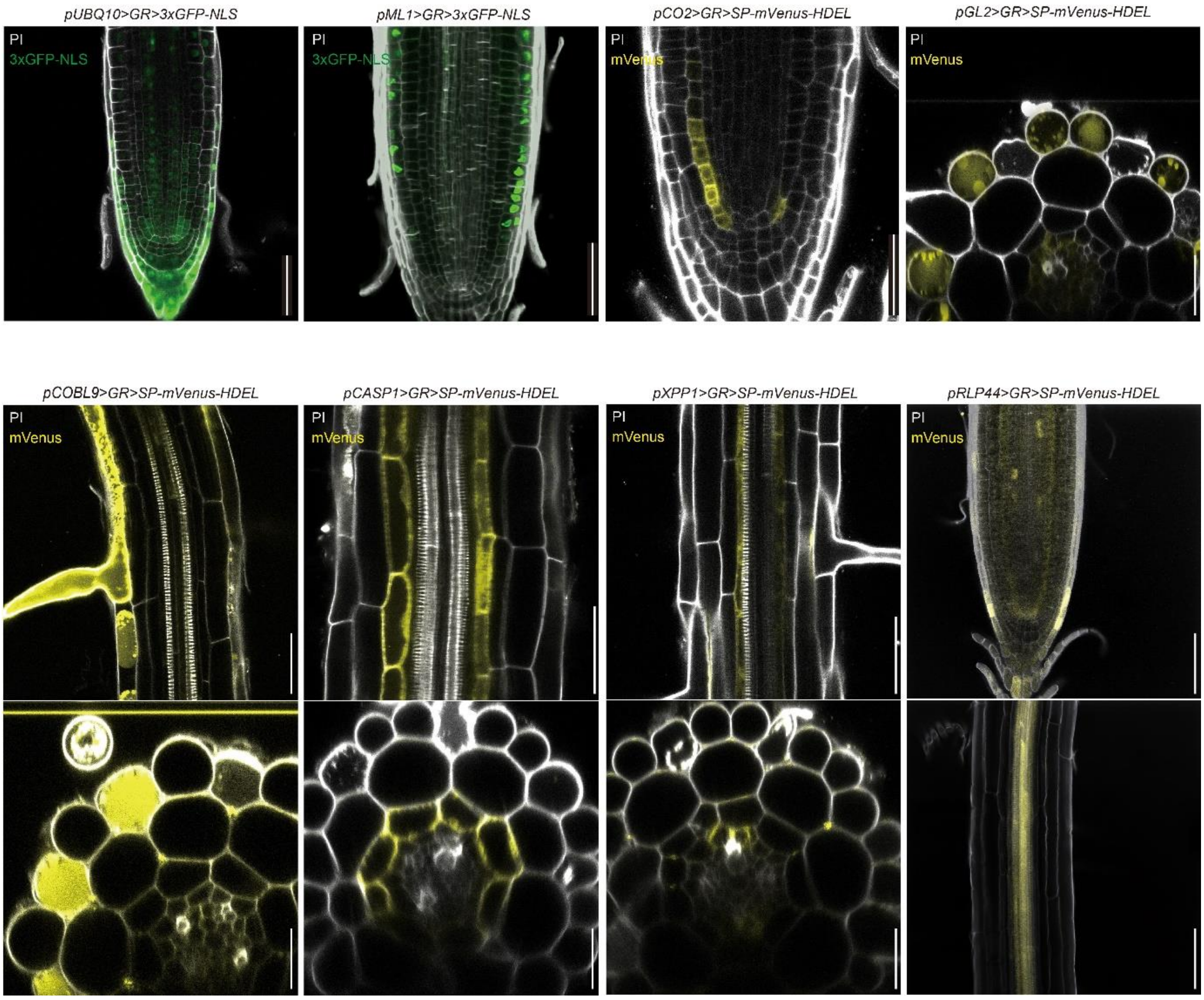

Supplemental Figure 3. Fluorescent reporters under control of the pOp promoter demonstrating tissue-specific expression in –GR-LhG4 trans-activation lines. Fluorescence derived from the indicated reporter (pOp6:3xGFP-NLS or pOp6:SP-mVenus-HDEL) in lines expressing the GR-LhG4 trnascription factor in the epidermis (under control of the *pML1* promoter), in meristematic cortex cells *(pCO2*), in trichoblasts (*pCOBL9),* atrichoblasts (*pGL2*), in differentiating and mature endodermis cells (*pCASP1)*, in xylem pole pericycle cells (*pXPP)*, and in most RAM tissues with highest expression in the vascular tissue under control of the *pRLP44* promoter. Cells walls are counter-stained with PI. Bars = 50 µm in longitudinal sections and = 20 µm in cross sections. Cross sections are from the differentiation zone of the root before formation of the casparian strip.

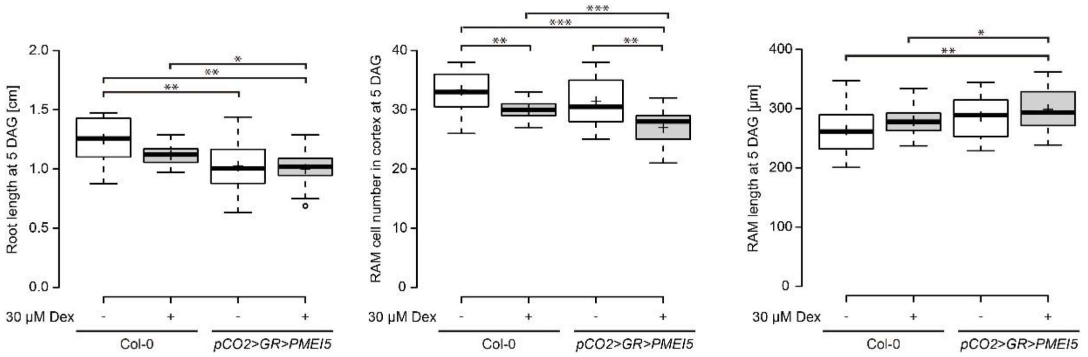

Supplemental Figure 4. Expression of *PMEI5* in meristematic cortex cells (pCO2>GR>PMEI5) had only mild effect on primary root growth and RAM morphology. (A) Primary root length (A), RAM cell number (B), and RAM length (C) five days after germination in Col-0 and pCO2>GR>PMEI5 roots in the absence and presence of the inducer dexamethasone (Dex). For (B) and (C) n = 40 - 50. Asteriks indicate statistically significant difference revealed by a two-tail t-test with $p < 0.05$ (*), $p < 0.01$ (**) and $p < 0.001$ (***).

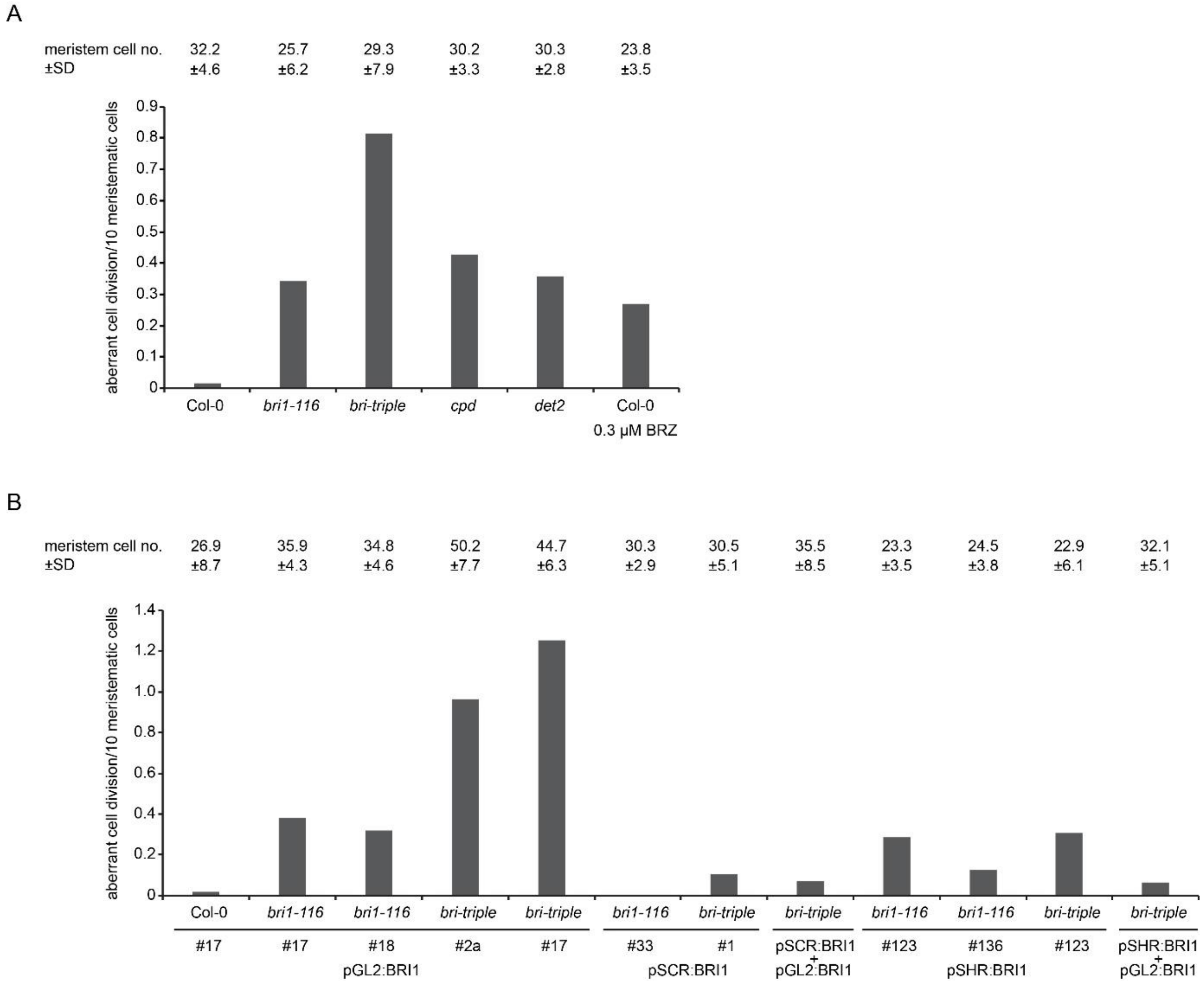

Supplemental Figure 5. Aberrant cell divisions in meristems of BR mutants and plants grown under BR-limiting conditions, corresponding to Figure 5. Bars show number of aberrant cell divisions in the meristems of the indicated genotypes per 10 meristematic cortex cells. Numbers above bars indicate average meristematic cell number ±S.D. Analysis was performed on samples used for Figure 5.

**Table S1:** Mutants and transgenic lines used in this study

| Mutant/transgenic line | Reference | Accession |
|---|---|---|
| *PMEIox* | Wolf et al., 2012 | |
| *cnu1* | Wolf et al., 2012 | |
| *cnu2* | Wolf et al., 2014 | |
| *pUBQ10>GR>PMEI5* | this study | |
| *pML1>GR>PMEI5* | this study | |
| *pCOBL9>GR>PMEI5* | this study | |
| *pGL2>GR>PMEI5* | this study | |
| *pCO2>GR>PMEI5* | this study | |
| *pCASP1>GR>PMEI5* | this study | |
| *pXPP>GR>PMEI5* | this study | |
| *pRLP44>GR>PMEI5* | this study | |
| *bri1-116* | Li and Chory 1997 | |
| *bri triple* | (Vragovic et al., 2015) | *bri1-116*;<br>SALK_005982 (brl1);<br>SALK_006024 (brl3) |
| *pGL2:BRI1-GFP* | Fridman et al., 2014 | |
| *pGL2:BRI1-GFP (bri1-116)* | Hacham et al., 2011 | |
| *pGL2:BRI1-GFP (bri-triple)* | Vragovic et al., 2015 | |
| *pSCR:BRI1-GFP (bri1-116)* | Hacham et al., 2011 | |
| *pSCR:BRI1-GFP (bri-triple)* | Vragovic et al., 2015 | |
| *pSHR:BRI1-GFP (bri1-116)* | Hacham et al., 2011 | |
| *pSHR:BRI1-GFP (bri-triple)* | Vragovic et al., 2015 | |
| *cpd* | Szekeres et al., 1996 | |
| *det2* | Chory et al., 1991 | |
| *GFP-TUA6* | Ueda et al., 1999 | |
| *GFP-LTI6b/H2B-RFP* | *Maizel et al., 2011* | |
| *35S:RLP44-GFP (cnu2)* | Wolf et al., 2014 | |

| | | | |
|---|---|---|---|
| **pSW507** | pUBQ10 intermediate | | |
| pSW336 | pUBQ10 (pGGA0006) | Lampropoulos et al., 2013 | |
| pSW182 | B-Dummy (pGGA022) | Lampropoulos et al., 2013 | |
| pSW181 | GR-LhG4 | Schürholz et al., 2018 | |
| pSW184 | D-dummy (pGGD002) | Lampropoulos et al., 2013 | |
| pSW185 | Rbcs terminator (pGGE001) | Lampropoulos et al., 2013 | |
| pSW188 | F-H adapter | Lampropoulos et al., 2013 | |
| pGGM000 | Intermediate vector | Lampropoulos et al., 2013 | |
| **pSW304** | pML1 intermediate | | |
| pSW179 | pML1 (pGGA0022) | Schürholz et al., 2018 | |
| pSW182 | B-Dummy (pGGA022) | Lampropoulos et al., 2013 | |
| pSW181 | GR-LhG4 | Schürholz et al., 2018 | |
| pSW184 | D-dummy (pGGD002) | Lampropoulos et al., 2013 | |
| pSW185 | Rbcs terminator (pGGE001) | Lampropoulos et al., 2013 | |
| pSW188 | F-H adapter | Lampropoulos et al., 2013 | |
| pGGM000 | Intermediate vector | Lampropoulos et al., 2013 | |
| **pSW306** | pCOBL9 intermediate | | |
| pSW222 | pCOBL9 | AACAGGTCTCAACCTTATGTACTCTTTTAGTGGTTTACAC | AACAGGTCTCATGTTTGTGTCTTTCTCCAGAGAAAG |
| pSW182 | B-Dummy (pGGA022) | Lampropoulos et al., 2013 | |
| pSW181 | GR-LhG4 | Schürholz et al., 2018 | |
| pSW184 | D-dummy (pGGD002) | Lampropoulos et al., 2013 | |
| pSW185 | Rbcs terminator (pGGE001) | Lampropoulos et al., 2013 | |
| pSW188 | F-H adapter | Lampropoulos et al., 2013 | |
| pGGM000 | Intermediate vector | Lampropoulos et al., 2013 | |
| **pSW307** | pGL2 intermediate | | |

| | | | |
|---|---|---|---|
| pSW223 | pGL2 | AACAGGTCTCAACCTCGTATTATACGGACGGTTTAAGC | AACAGGTCTCATGTTTTTTCTTCTTAATATTCGATTTTTAATA |
| pSW182 | B-Dummy (pGGA022) | Lampropoulos et al., 2013 | |
| pSW181 | GR-LhG4 | Schürholz et al., 2018 | |
| pSW184 | D-dummy (pGGD002) | Lampropoulos et al., 2013 | |
| pSW185 | Rbcs terminator (pGGE001) | Lampropoulos et al., 2013 | |
| pSW188 | F-H adapter | Lampropoulos et al., 2013 | |
| pGGM000 | Intermediate vector | Lampropoulos et al., 2013 | |
| **pSW305** | pCO2 intermediate | | |
| pSW299 | pCO2 | AACAGGTCTCAACCTGATCAGAGTATTGGGCCTTTTGG | AACAGGTCTCATGTTTATCGTTATTAACTAGGGTTCTTG |
| pSW182 | B-Dummy (pGGA022) | Lampropoulos et al., 2013 | |
| pSW181 | GR-LhG4 | Schürholz et al., 2018 | |
| pSW184 | D-dummy (pGGD002) | Lampropoulos et al., 2013 | |
| pSW185 | Rbcs terminator (pGGE001) | Lampropoulos et al., 2013 | |
| pSW188 | F-H adapter | Lampropoulos et al., 2013 | |
| pGGM000 | Intermediate vector | Lampropoulos et al., 2013 | |
| **pSW302** | pCASP1 intermediate | | |
| pSW177 | pCASP1 | Schürholz et al., 2018 | |
| pSW182 | B-Dummy (pGGA022) | Lampropoulos et al., 2013 | |
| pSW181 | GR-LhG4 | Schürholz et al., 2018 | |
| pSW184 | D-dummy (pGGD002) | Lampropoulos et al., 2013 | |
| pSW185 | Rbcs terminator (pGGE001) | Lampropoulos et al., 2013 | |
| pSW188 | F-H adapter | Lampropoulos et al., 2013 | |
| pGGM000 | Intermediate vector | Lampropoulos et al., 2013 | |
| **pSW303** | pXPP intermediate | | |
| pSW178 | pXPP | Schürholz et al., 2018 | |
| pSW182 | B-Dummy (pGGA022) | Lampropoulos et al., 2013 | |
| pSW181 | GR-LhG4 | Schürholz et al., 2018 | |

| | | | |
|---|---|---|---|
| pSW184 | D-dummy (pGGD002) | Lampropoulos et al., 2013 | |
| pSW185 | Rbcs terminator (pGGE001) | Lampropoulos et al., 2013 | |
| pSW188 | F-H adapter | Lampropoulos et al., 2013 | |
| pGGM000 | Intermediate vector | Lampropoulos et al., 2013 | |
| **pSW308** | pRLP44 intermediate | | |
| pSW299 | pRLP44 | AACAGGTCTCAACCTTTTGCGATATTTTTGGCTGTC | AACAGGTCTCATGTTTTTTAAATTTTAGAGAGGTTTC |
| pSW182 | B-Dummy (pGGA022) | Lampropoulos et al., 2013 | |
| pSW181 | GR-LhG4 | Schürholz et al., 2018 | |
| pSW184 | D-dummy (pGGD002) | Lampropoulos et al., 2013 | |
| pSW185 | Rbcs terminator (pGGE001) | Lampropoulos et al., 2013 | |
| pSW188 | F-H adapter | Lampropoulos et al., 2013 | |
| pGGM000 | Intermediate vector | Lampropoulos et al., 2013 | |
| **pSW301** | PMEI5 intermediate | | |
| pSW189 | H-A adapter | Lampropoulos et al., 2013 | |
| pSW180 | pOp6 (pGGA016) | Lampropoulos et al., 2013 | |
| pSW182 | B-Dummy (pGGA022) | Lampropoulos et al., 2013 | |
| pSW190 | PMEI5 | AACAGGTCTCAGGCTCAATGGCCACAATGCTAATAAACCAC | AACAGGTCTCACTGATTACTTATTTTCAACAAGCTTGTGACC |
| pSW184 | D-dummy (pGGB003) | Lampropoulos et al., 2013 | |
| pSW186 | UBQ10 terminator (pGGE009) | Lampropoulos et al., 2013 | |
| pSW187 | KanR (pGGF007) | Lampropoulos et al., 2013 | |
| pGGN000 | Intermediate vector | Lampropoulos et al., 2013 | |

**Table S2**. Overview of constructs generated with GreenGate cloning (Lampropoulos et al., 2013) and the primers used to generate modules, where appropriate.
